# DORA: an interactive map for the visualization and analysis of ancient human DNA and associated data

**DOI:** 10.1101/2024.01.15.575663

**Authors:** Keith D. Harris, Gili Greenbaum

## Abstract

The ability to sequence ancient genomes has revolutionized the way in which we study evolutionary history by providing access to the most important aspect of evolution — time. Until recently, studying human demography, ecology, biology, and history using population genomic inference relied on contemporary genomic datasets. Over the past decade, the availability of human ancient DNA (aDNA) has increased rapidly, almost doubling every year, opening the way for spatiotemporal studies of ancient human populations. However, the multidimensionality of aDNA, with genotypes having temporal, spatial and genomic coordinates, and the need to integrate multiple sources of data, poses a challenge for developing meta-analyses pipelines. To address this challenge, we developed a publicly-available interactive tool, `DORA`, which integrates multiple data types, genomic and non-genomic, in a unified interface. This web-based tool allows users to browse sample metadata along with additional layers of information, such as population structure, climatic data, and unpublished samples. Users can then perform analyses on genotypes of these samples, or export sample subsets for external analyses. `DORA` integrates analyses and visualizations in a single intuitive interface, resolving the technical issues of combining datasets from different sources and formats, and allowing researchers to focus on analysis and the scientific questions that can be addressed through analysis of aDNA datasets.

## 1 Introduction

Until recently, answering questions regarding evolution and demographics of human history has relied on inference from contemporary sequenced genomes and a small number of sequenced ancient genomes [1]. Over the past decade, the availability of human ancient DNA (aDNA) has increased rapidly, almost doubling every year [2]. This data includes high-quality whole genome shotgun sequencing for a very small number of samples of interest, low coverage whole genome shotgun sequencing, and sequence capture targeting SNPs in specific SNP panels (e.g., Human Origins, 1240K [3, 4]). These resources now allow for testing evolutionary questions directly, rather than relying on indirect inference, by tracking changes in allele frequencies, genomic compositions, and phenotypic predictions over many millennia [5–8].

The visualization and analysis of aDNA is challenging, partly owing to the multi-dimensionality of the individual data points. For every genotype from an individual sample in the database, there is a (i) genomic position, (ii) geographic location, and (iii) temporal position (sample dating, which also varies between dating methods). Different analyses can consider a subset of genotypes along windows in each of these dimensions. Selection of such windows can be based on geographic regions of interest, time periods of interest, and also on accessory data such as sample quality, specified regions of the genome, climatic databases, geographical boundaries, and other considerations relevant to the specific research question. One of the challenges of meta-analyses of human ancient genomics is the integration and visualization of these multiple sources of data, which is necessary to allow selection of the specific genotypes to be analyzed. Such selection procedures are often informed by exploratory analyses of the dataset. Tools have been developed that are specifically designed to visualize the results of analyses of aDNA [9], but these require additional tools in order to generate these results. Ideally, the first steps of such meta-analyses would be conducted within a single framework that would allow exploration of the dataset by allowing intuitive selection of spatial, temporal, and genomic windows of genotypes, along with population genetic analyses of the selected genotypes.

Here, we present an interactive map of human aDNA samples, called **D**ata **O**verlays for **R**esearch in **A**rchaeogenomics (`DORA`). This tool visualizes aDNA samples on a geographic map, and allows selection of samples by visualizing the characteristics of the samples (e.g., metadata of the sample such as coverage). It also includes additional data layers that can be displayed alongside the samples, such as climatic data and results of downstream population structure analyses. Importantly, the tool is designed to allow users to integrate these data layers in order to select subsets of samples and genotypes according to regions that can be drawn on the map, and windows that can be selected using the slider under the map. These subsets can be exported, or used in meta-analyses through the `DORA` interface. The interface is web-based and its resources are stored on Amazon Web Services Simple Storage Service (AWS S3), while private user data added to the tool will be available only on the local machine and not uploaded to AWS S3. `DORA` can facilitate a range of meta-analyses of ancient genomes and will allow users to seamlessly integrate published resources with their unpublished genomic samples or other data layers.

To demonstrate the usefulness of this tool, we have pre-loaded a curated aDNA dataset, the *Allen Ancient DNA Resource* (AADR) [2], along with the TraCE21K climatic data from the database *Climatologies at high resolution for the earth’s land surface areas* (CHELSA) [10]. We also developed an interface for loading polygenic scores (PGSs), which can be used to predict the phenotypes of the ancient samples based on their genomes, from the PGSCatalog database [11]; `DORA` is designed to directly apply PGSs to the aDNA samples loaded into the tool. We demonstrate some of the analyses that can be performed with the tool, including tracking allele frequencies in geographically defined populations, computing temporal trends in genetic differentiation between populations using pairwise *F*_*ST*_ and PCA, and tracking polygenic score values of individuals and demarcated populations over time. `DORA` integrates this variety of data types, analyses and visualizations, in a single intuitive interface, resolving the technical issues of combining datasets from different sources and formats, and allowing researchers to focus on the analysis and the scientific questions of interest.

## 2 Features of DORA

### 2.1 Map interface

The main display of `DORA` is a world map on top of which data is visualized. By default, `DORA` loads a topographical map of the world in equirectangular projection [12] and sample metadata (a screenshot of the main display with metadata loaded from AADR is shown in Figure 1). Samples are represented as dots on the map, with the color of the dots reflecting additional sample metadata, such as coverage or dating, as defined in the ‘Colormap’ box on the right. The map can be browsed spatially by zooming into specific regions and resetting by clicking ‘Reset view’ after right-clicking the map. At the bottom of the map is a timeline that shows the number of available samples in predefined temporal bins (in 100s of years). A slider on the timeline with a blue transparent background defines which samples will be displayed on the map; users can shift the sides of the slider to define this window. In addition, a ‘variants panel’ can be opened by clicking on the ‘Variants’ tab at the bottom right of the screen. This displays selected variants according to their genomic regions; new regions can be added by clicking anywhere in the panel, while selected regions can be removed by clicking on the yellow box representing the region. Variant selection in this panel is used for analyses that are run on multiple loci, such as *F*_*ST*_ or PCA (see the *Example analyses* section below).

**Figure 1:**
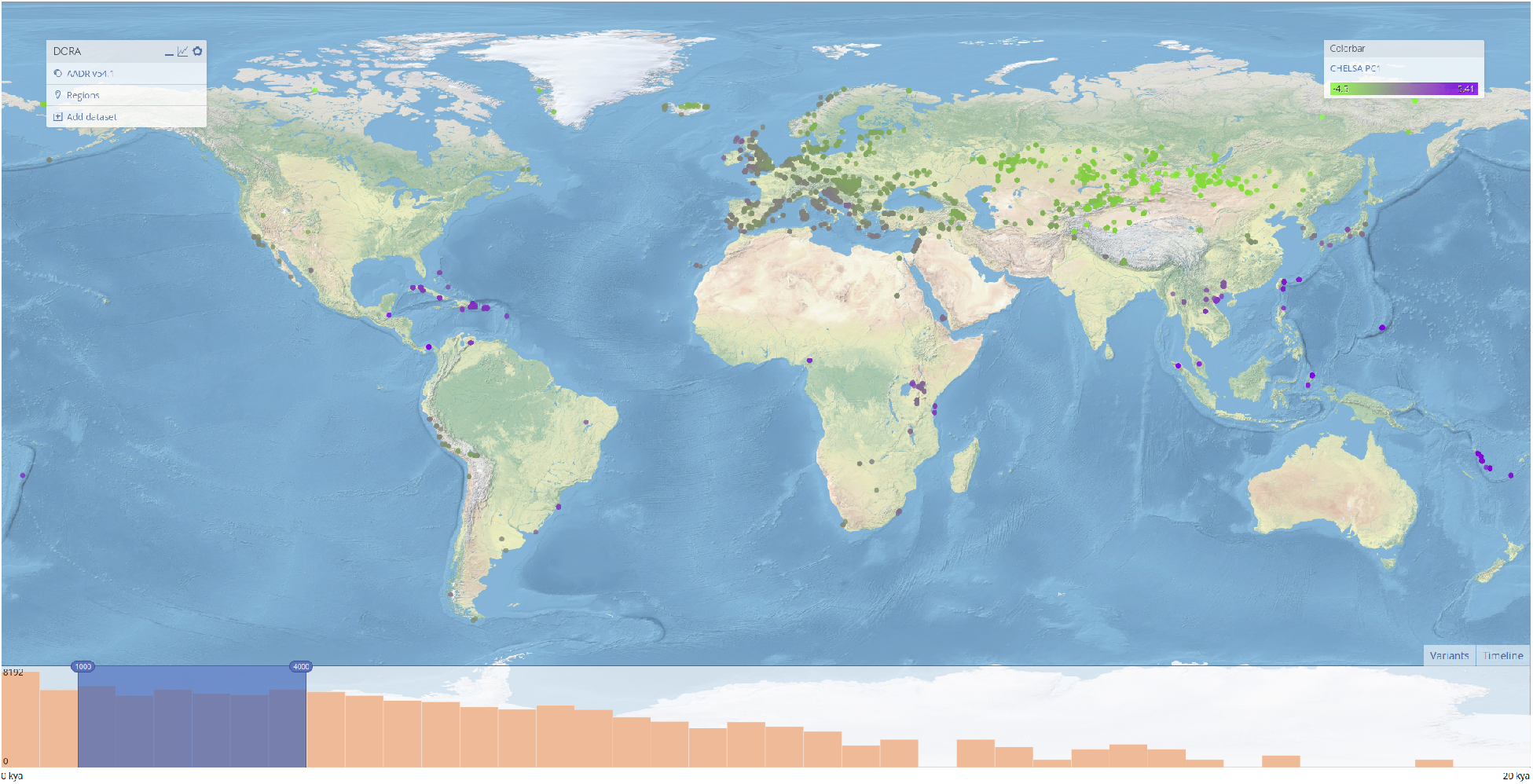
Main interface of DORA. Layers are shown in the main menu on the top left corner. Samples appear as dots on the map, with the color reflecting a specific attribute of the sample, that can be selected on the ‘Colorbar’ panel on the right. In this example, we selected the ‘CHELSA PC1’ attribute, which reflects predicted climate differences between samples based on their geolocation and dating. At the bottom of the map is a timeline of sample data within the displayed geographic boundaries, and the variants panel, that can be opened by clicking the ‘Variants’ tab. The blue highlighted bins indicate the year range of displayed samples and can be moved to select specific ranges.

To select samples according to their geographic positions, users can define ‘Regions’, which are colored polygons that demarcate samples for subsequent analyses or export. Regions are created by right-clicking anywhere on the map and clicking ‘Create Region’. Once clicked, the next successive left-clicks on the map will create vertices of the polygon. Right-clicking will save the polygon with the marked vertices. Regions are stored in the browser cache until deleted; regions can be deleted by clicking a region and pressing the backspace or delete key. Double-clicking on a region will allow the user to edit the region label, or exclude samples in the region from analyses.

### 2.2 Adding datasets

`DORA` overlays multiple data sources onto a single map display, allowing users to combine these sources in selected samples for subsequent analyses (Fig. 2). Layers that were pre-loaded to demonstrate the features of `DORA` will load by default, including the map (based on Natural Earth [12]), aDNA metadata and genotypes from the AADR, and CHELSA climate model data for the past 20,000 years. Additional datasets can be added that either add samples to the map (primary datasets, see the scheme in Figure 3), or add metadata for existing samples, according to their sample IDs. If samples have multiple possible attributes to display, the user is able to select the desired attribute to color the samples using the ‘Colormap’ menu near the top right corner of the map.

**Figure 2:**
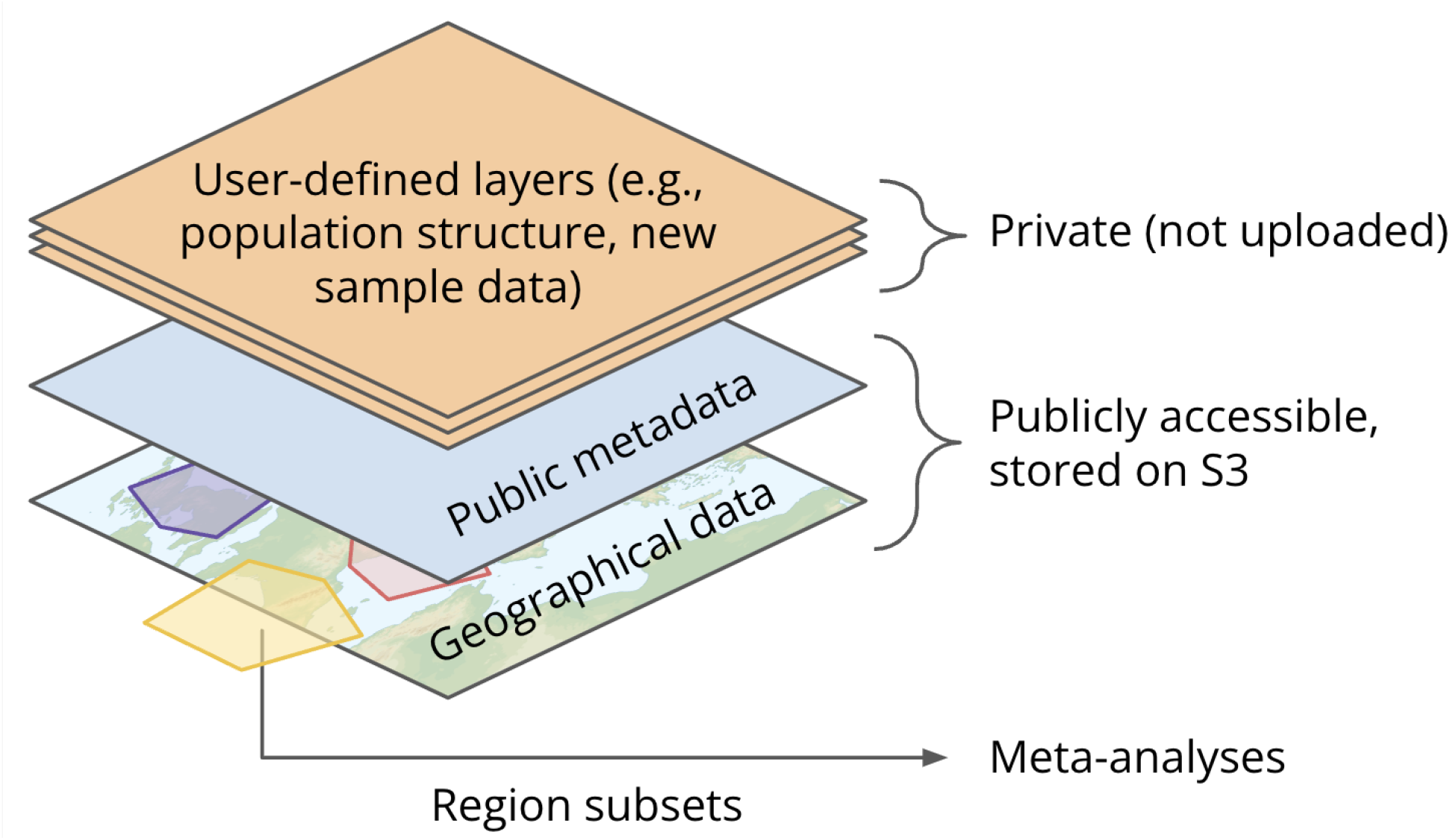
Schematic representation of the organization of data layers in DORA. Pre-loaded data includes publicly available datasets, including the AADR and CHELSA TraCE21K. Additional user-defined layers can be added by users, and will be accessible only on the local machine of the user. Analyses can be conducted on all available data.

**Figure 3:**
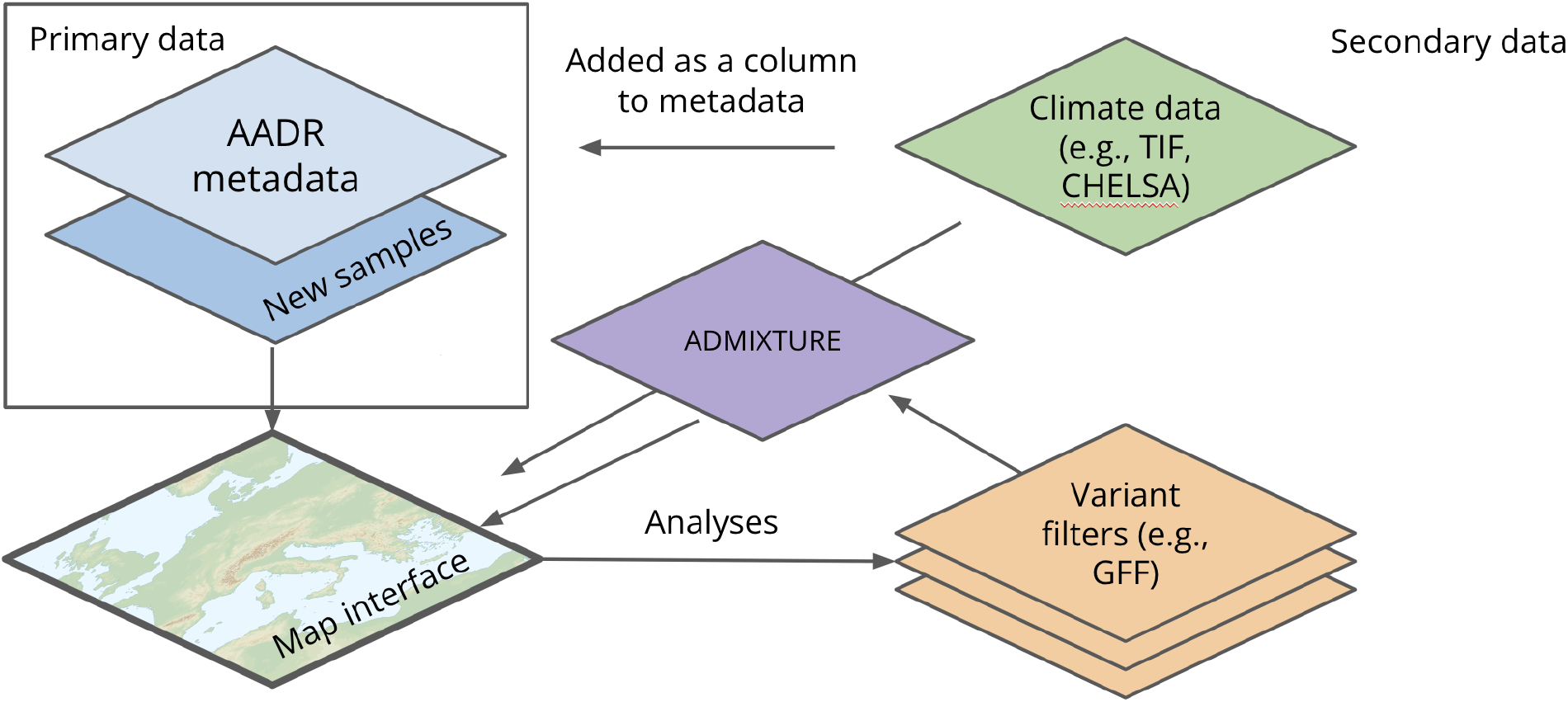
Schematic representation of how different layers are integrated in DORA. Primary data includes files that provide mandatory metadata of samples, including a date estimate and geolocation. Accessory metadata can be added from a variety of sources, such as climate data for sample locations or ADMIXTURE results. This data can then be displayed on the map, used to filter individuals for analyses, or plotted along with the results of genetic analyses.

Additional datasets can be added by clicking the ‘Add Dataset’ link on the main menu; a schematic of the integration of different types of data is presented in Figure 3. Previously added datasets will appear in the ‘Library’ tab of the ‘Add Dataset’ dialog box. `DORA` supports the addition of sample metadata for existing or additional samples (.csv/.tsv files), genotypes for samples with loaded metadata that can be used for analyses (PLINK .bed/.bim/.fam files), ADMIXTURE results (.fam/.Q files), and PGSs loaded from PGSCatalog through the tool interface (Table 1). User-defined layers and settings will persist in accordance with browser cache retention, which varies between browsers.

**Table 1:**
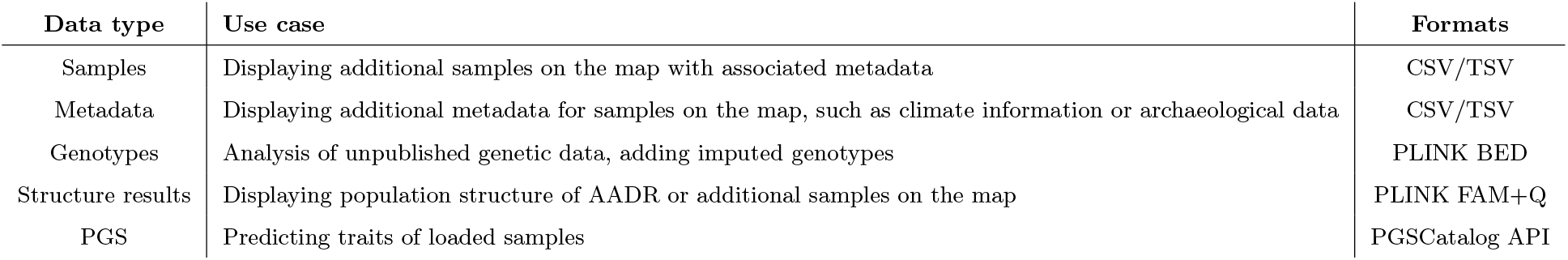
Data types supported by DORA.

| Data type | Use case | Formats |
| --- | --- | --- |
| Samples | Displaying additional samples on the map with associated metadata | CSV/TSV |
| Metadata | Displaying additional metadata for samples on the map, such as climate information or archaeological data | CSV/TSV |
| Genotypes | Analysis of unpublished genetic data, adding imputed genotypes | PLINK BED |
| Structure results | Displaying population structure of AADR or additional samples on the map | PLINK FAM+Q |
| PGS | Predicting traits of loaded samples | PGSCatalog API |

### 2.3 Analyses

`DORA` allows users to conduct meta-analyses on the genetic data loaded into the interface, which includes the AADR genotypes loaded into `DORA` by default, and any other data loaded into the browser (in PLINK BED format). One of the main benefits of running analyses from the map interface is that users can integrate multiple data sources when defining subsets of samples, and run a variety of analyses on these subsets from the same interface. Analyses can be initiated by right-clicking anywhere on the map and clicking ‘Analyze’. This will open a dialog box with tabs for available analyses. Analyses are run on the selected geographic regions, in temporal windows assigned according to the timeline or in the analysis dialog box field ‘Year range’, and on genomic regions as defined in the variants panel (except the allele frequency analysis, which runs on a single variant specified in the anlaysis dialog box). The available analyses include: (i) allele frequency analysis, which shows the temporal trajectory of the allele frequency of a specific variant; (ii) expected heterozygosity of selected genetic variants; (iii) pairwise *F*_*st*_ between all the pairs of regions (using the implementation of Hudson *F*_*st*_ estimation [13] from `scikit-allel` [14]); (iv) PCA of samples from all regions; and (v) computed polygenic scores of samples in all regions, over the chosen temporal windows (PGSs must first be imported from `PGSCatalog`).

Analyses are run on AWS Lambda for publicly available data, and on the local machine for locally stored data. `DORA` uses `Pyodide` [15] to provide the same Python analysis environment in the browser and on AWS Lambda. The analysis script is loaded by the specific environment, and the BED file is read either from AWS S3 (in the case of AWS Lambda) or from the browser cache (in the case of local data). The user receives a link to the analysis results when the analysis is run; this link will display the results as soon as they are ready. When the analysis is complete, the results are also processed and plotted using Python code run using `Pyodide`. Each analysis comes with a number of pre-defined plotting formats, such as line plots or scatter plots; this Python code can be manipulated by the user, and the figure updated using the ‘Update’ button (Fig. 4). The use of the same Python environment for both analysis and result plotting allows for flexibility and accessibility of the analysis and plotting code, and supports straightforward development of additional analyses. The outputs of the analyses are broad enough to allow many possible visualizations using the same analysis, by modifying the plotting code. All analysis plots presented here were generated by the analysis feature of `DORA`.

**Figure 4:**
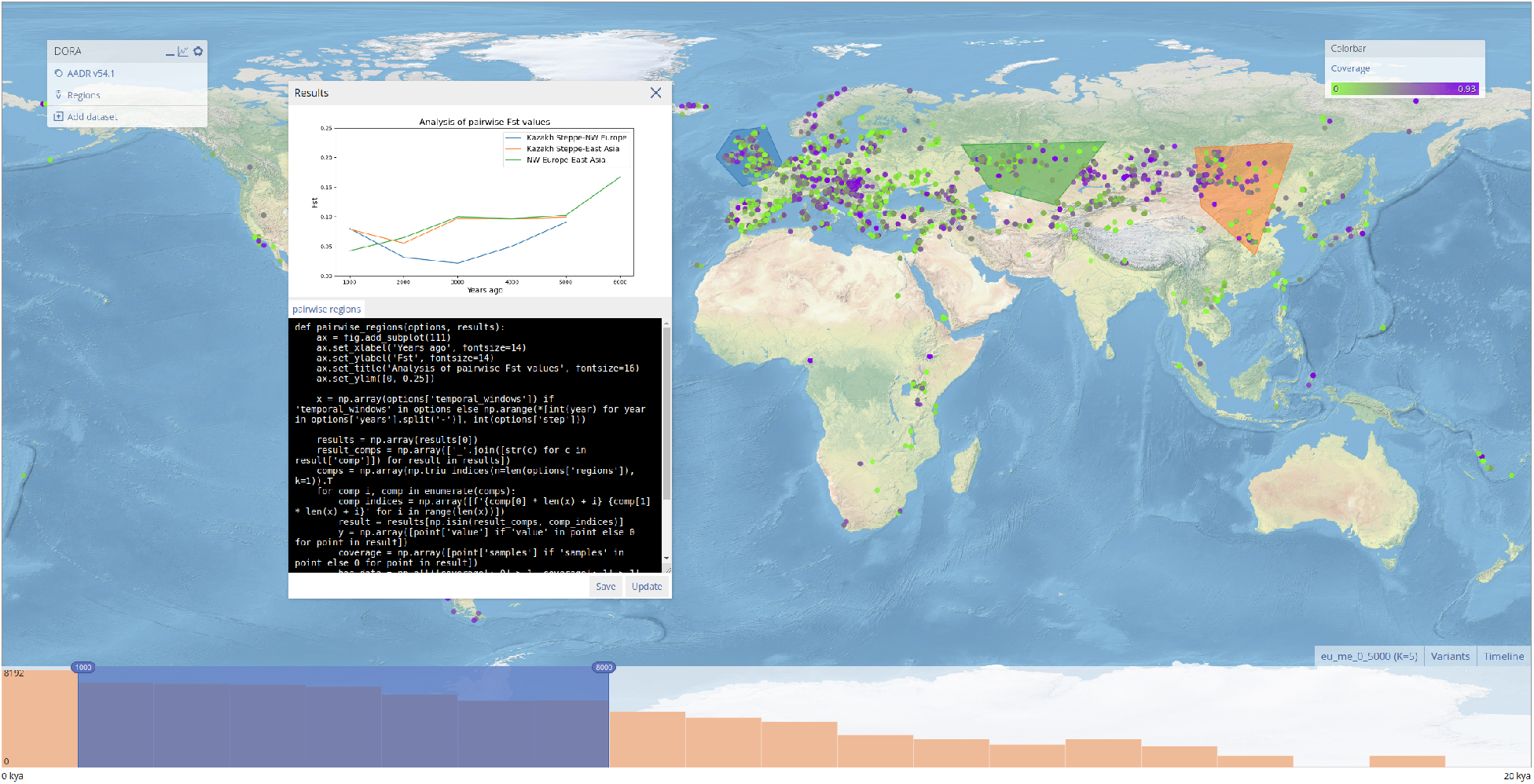
Display of analysis results in DORA. When an analysis is completed, the results open as a dialog box with a Python-generated plot above a code box (here shown to the left of the selected regions). In this example, the results of a pairwise *F*_*ST*_ analysis of selected regions are plotted (for details, see the *Example analyses* section below). Users can edit the code and update the plot by clicking ‘Update’, or download the plot by clicking ‘Save’. Previous results will also be available in the list of previous analyses that can be opened by clicking the graph icon in the top right corner of the main menu.

In the analysis dialog box, users can define filtering criteria for samples and variants, including (i) options specific to the analysis type, (ii) filtering that relates to metadata of the samples, and (iii) the proportion of missing genotypic data for the samples and variants.

## 3 Example analyses

To illustrate some potential uses of `DORA`, we present here example analyses that utilize different features of the tool. These tasks did not require the preparation of data with external tools and analyses were completed in less than one minute. The results are also processed and plotted by `DORA`, so that the entire task is completed using the same interface.

### 3.1 Region selection based on ADMIXTURE analysis

One of the basic functions of the map is the selection of geographic regions, which can be used to define subsets of individuals that will be grouped in subsequent analyses. As mentioned above, regions are created dynamically by drawing polygons on the map. In this example, we wanted to use a previously conducted population structure analysis, conducted with ADMIXTURE [16]. The ADMIXTURE results are represented in Figure 5 as colored samples on the map, where each individual is colored by its main ancestry component. At the bottom of the map, above the timeline, a ‘STRUCTURE-plot’ is shown for all displayed samples. Based on these results, we drew three polygons representing Northwest Europe, the Kazakh Steppe and East Asia.

**Figure 5:**
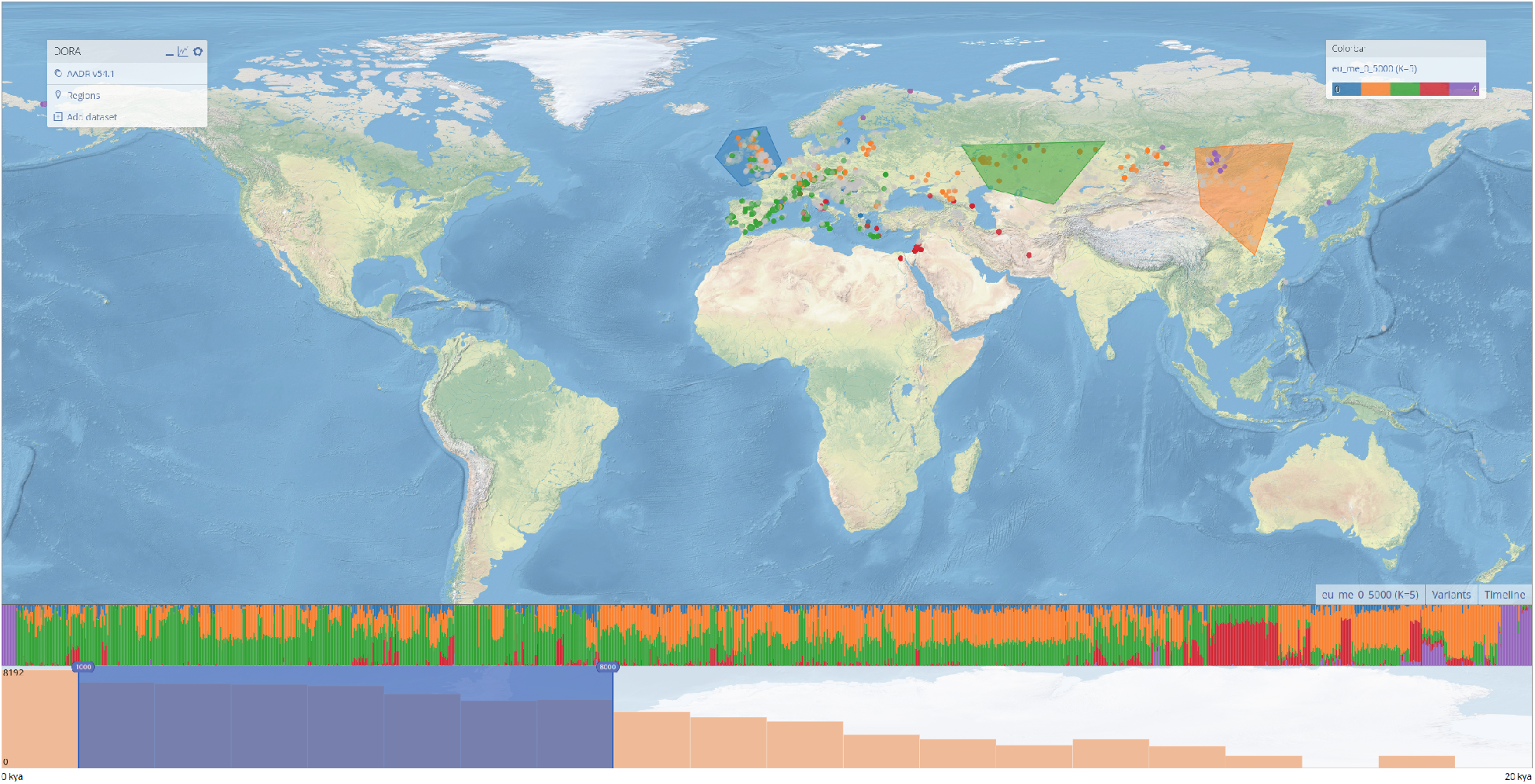
Selection of regions based on population structure analysis. ADMIXTURE results are plotted on the map, with samples colored according to their largest ancestry component. This can be selected using the ‘Colorbar’ menu on the top right, after adding the ADMIXTURE results (see *Adding datasets*). The STRUCTURE-plot for all samples in the selected window is presented above the timeline, ordered according to longitude. Regions have been manually selected on the map to roughly correspond to the plotted ADMIXTURE results.

### 3.2 Allele frequency fluctuations over time in different populations

Once regions are selected (in this example, with the aid of the integrated ADMIXTURE results), downstream analyses can be run. By default, a subset of variants in the data is used for analyses; this can be changed in the variants panel at the bottom of the screen. Note that increasing the number of variants will increase the running time of the analysis, and if the number of variants is too large some analyses may fail to run.

A straightforward and common analysis that can be performed is tracking allele frequencies over time in the selected geographic and temporal windows. We selected two variants that are at intermediate frequencies in modern populations: a variant that was suggested to have undergone positive selection during the Black Plague [8, 17] (`rs2549794`), and a randomly selected variant (`rs262680`). In the windows selected, the frequency of the non-reference allele of the `rs262680` variant increases monotonically in Northwest Europe and fluctuates in the other regions (Fig. 6A), whereas `rs2549794` shows no clear trend in any region (Fig. 6B). Such analyses may be useful to examine whether observed trends are common or if they indeed reflect a selection process.

**Figure 6:**
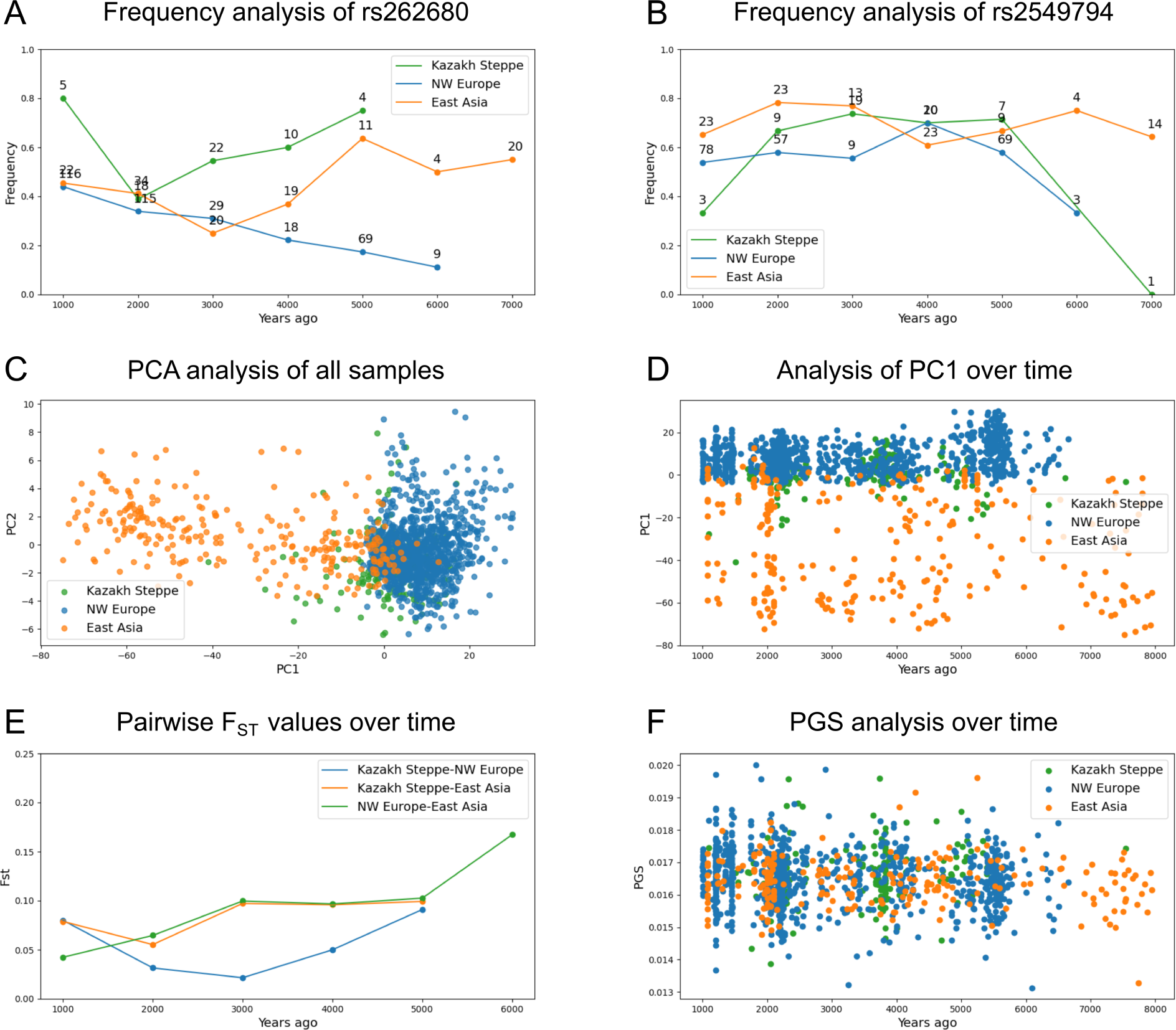
Example analyses run with DORA. Using the region selection tool, we defined three geographic regions (Northwest Europe in blue, East Asia in orange, and the Kazakh Steppe in green). We ran different analyses on these regions. (**A**) Temporal changes in allele frequency of variant `rs262680`. (**B**) Temporal changes in allele frequency of variant `rs2549794`. (**C)** PCA of samples from all temporal windows in all three regions for a subset of variants from chromosome 1. (**D**) PC1 of the three regions over time using the same PCA from panel (**C**). (**E**) Pairwise *F*_*ST*_ values between the three regions over time, using the same subset of variants from panel (**C**). (**F**) Height PGS (computed using data integrated from the PGSCatalog) for samples in all three regions over time (PGSCatalog ID `PGS000297`).

### 3.3 PCA of samples in selected regions

`DORA` can run PCA on a subset of samples, for specified genomic regions. By default, `DORA` pools samples from all windows and runs the PCA on all selected variants. To deal with missing data, `DORA` includes a few imputation approaches that can be selected in the analysis dialog box (e.g., sampling genotypes using the allele frequency of all samples or of the subset, or setting the frequency as 0). We show here two possible displays of this result: a scatter plot of the first two principal components (Fig. 6C), and a scatter plot of the temporal position of the samples and the first principal component (Fig. 6D).

In the standard PCA plot, region selection captures genetic differences between the samples located in these regions (Fig. 6C). Overlap can be seen mainly between the Kazakh Steppe and Northwest Europe, but also between East Asia and the Kazakh Steppe. To see whether this overlap occurs in specific temporal windows, we can plot the dating of individual samples against the first principal component: here the separation of samples from different regions is apparent in each selected period, with increasing overlap between East Asia and Northwest Europe closer to the present (Fig. 6D).

### 3.4 Computing pairwise *F*_*st*_ values for region pairs

We can also visualize genetic differentiation at a population-level using the *F*_*ST*_ statistic. `DORA` can compute pairwise *F*_*ST*_ values for groups of samples defined by region, temporal window, and subsets of genetic variants. As an analog to (Fig. 6D), we computed pairwise *F*_*ST*_ values for pairs of regions within each temporal window (Fig. 6E).

The results show high genetic differentiation between the three regions that decreases towards the present, with a drastic decrease in *F*_*ST*_ around the period of the Yamnaya expansion (4000-5000ya) [18]. It is important to note that *F*_*st*_ values are also affected by the aspects of the scenario such as the number of samples used to calculate the *F*_*st*_ value in each window, population structure, and within-population variance [19], which may also affect the *F*_*st*_ values. The higher *F*_*ST*_ value for the Kazakh Steppe pairwise values in the 1000-2000ya temporal window can most likely be attributed to different sample sizes in this window. Such pairwise *F*_*st*_ analyses can be used together with population structure analyses as a means to define distinct regions that show genetic differentiation, before running downstream analyses.

### 3.5 Applying polygenic scores (PGSs) to aDNA samples

Polygenic scores (PGSs) are used to compute predicted trait values for an individual by weighting variants according to their contribution to the trait. Applying PGSs that have been developed using contemporary GWAS to aDNA samples provides an interesting opportunity to predict phenotypically meaningful genetic changes beyond single variants. In some cases, predicted traits can be correlated with measurable phenotypes, if they can be inferred from skeletal remains [7]. To allow crosstalk between the growing number of PGSs in the curated PGSCatalog [11], and aDNA samples, we added a feature to `DORA` that allows users to import PGSs directly from PGSCatalog, using a simple search interface. Users can search for a PGS according to the trait of interest, and the selected PGS will be processed to be compatible with genotypic data loaded into `DORA`.

To demonstrate this feature, we present the results of applying a PGS of height, which was developed based on the UK Biobank [20], to aDNA samples in the three regions. As with PCA results, the results of the PGS analysis can be displayed alongside other metadata of the samples, including their dating, coverage, climate, or additional user-defined metadata. We see no clear differences in the prediction of height between regions or temporal windows according to this PGS analysis (Fig. 6F). The application of PGSs to aDNA samples is not trivial due to the low transferability of PGSs between genetically distant populations [21]. Nevertheless, the exploration of temporal changes in predicted trait values for different traits in different regions of the world could provide an important lens through which human evolution can be studied; `DORA` not only simplifies conducting such analyses, it also allows integrating it with environmental data, potentially allowing for identification of drivers of such phenotypic changes.

## 4 Discussion

We present here `DORA`, a web-based tool for exploratory analysis of aDNA data. To demonstrate the features of this tool, we pre-loaded publicly available datasets and conducted a number of example analyses. These analyses were completed using the same interface and without using external tools. More elaborate analyses can be conducted with additional data layers.

The goal of `DORA` is to make the growing body of published human aDNA datasets more accessible for visualization and analysis, and to make the integration of accessory datasets for selection of samples and correlation with genetic analyses more intuitive. `DORA` combines serverless cloud-compute for analysis of publicly available datasets with local, browser-based analysis of private data. The results of these analyses are displayed in the browser, and their display can be manipulated by the user. Using the same Python environment for cloud-based and local computation, in addition to the generation of plots, makes the analysis pipelines on `DORA` transparent and accessible to users (Fig. 7).

**Figure 7:**
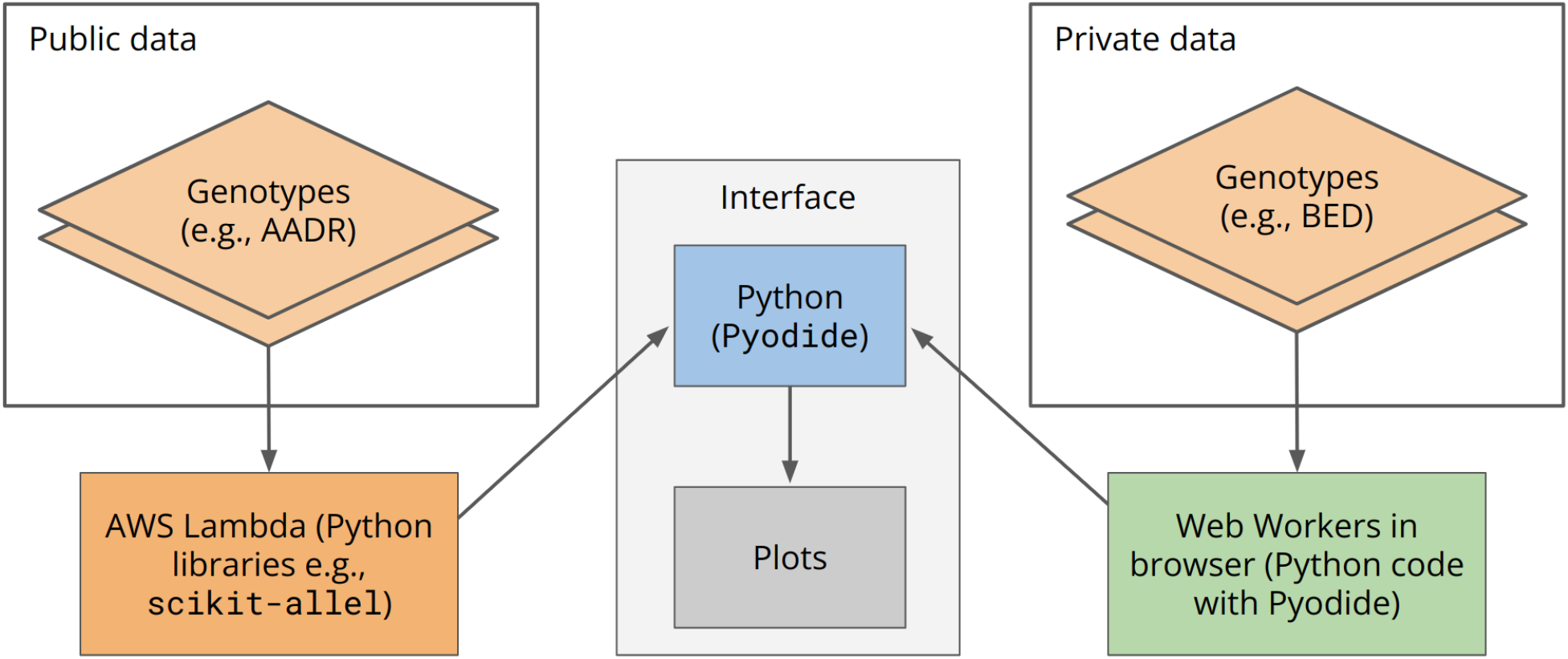
Schematic representation of how data is processed in different environments. Publicly available data is processed on cloud-based compute containers, such as AWS Lambda, while private data is processed in the browser. Both environments use the same Python analysis code, with the browser environment utilizing `Pyodide`. Results are then collected and plotted in the browser by Python code run with `Pyodide`.

Interface-based tools are particularly important for aDNA studies, because the multidimensionality of the data can pose a challenge to defining comprehensive and coherent criteria for selecting samples and genotypes for analyses. Downstream analyses also regularly involve multiple computational tools, each of which can have different requirements for the input format of the data. With currently available tools, such as PLINK and Python map plotting libraries, the visualization and analysis of aDNA involves multiple steps and libraries, and is not interactive; in `DORA`, these can be combined into a single step, and performed from the same interface.

We envision this tool as an important resource for researchers in the field of human aDNA by making accessible analyses that combine multiple existing resources with population genetics tools. This will allow rapid and intuitive definition of analyses and exploration of published aDNA data alongside unpublished samples.

## 5 Data availability

`DORA` is accessible at https://dora.modelrxiv.org. The latest version of the AADR is accessible at doi.org/10.7910/DVN/FFIDCW. The topographic map can be downloaded from Natural Earth [12]. The environmental variables used to compute the CHELSA PCs can be downloaded from the CHELSA website (https://chelsa-climate.org/chelsa-trace21k). PGSs used can be downloaded from PGSCatalog (https://www.pgscatalog.org).

## 6 Acknowledgments

We would like to thank Shai Carmi, David Gokhman, Viviane Slon, Fabrizio Mafessoni, Liran Carmel for comments, and members of the Greenbaum, Carmel and Gokhman labs for testing and commenting on features of the tool.

